# Consumption of the non-nutritive sweetener stevia for 12 weeks does not alter the composition of the gut microbiota

**DOI:** 10.1101/2022.11.17.516749

**Authors:** Gurdeep Singh, Andrew J McBain, John T McLaughlin, Nikoleta S Stamataki

## Abstract

The use of non-nutritive sweeteners (NNS) as an alternative to caloric sugars has increased in recent years. Stevia is a NNS that has demonstrated beneficial effects on appetite and energy intake. However, the impact on the gut microbiome is not well understood. Therefore, we investigated how regular consumption of stevia, for up to 12 weeks, impacts upon the human gut microbiome. Healthy subjects with a normal body mass index participated in the study; the stevia group (*n* = 14) were asked to consume five drops of stevia, twice daily, compared against control participants (*n* = 13). Faecal samples collected before and after treatment were analysed by 16S sequencing. Stevia did not cause significant changes in the beta or alpha diversity when compared to the control groups. When relative abundances of taxa were investigated, no clear differences could be detected. Conversely, random forest analysis could correctly associate the gut microbiome with control and stevia groups with an average of 75% accuracy, suggesting that there are intrinsic patterns that could discriminate between control and stevia use. However, large-scale changes in the gut microbiome were not apparent in this study and therefore, our data suggest that stevia does not significantly impact the gut microbiome.

## Introduction

Stevia is a non-nutritive sweetener (NNS) providing a sweet taste with no calories. Recent evidence supports a beneficial role of stevia and other NNS on energy intake and body weight [1,2]. More specifically, two randomised controlled trials from our group have shown beneficial effects related to the consumption of stevia-sweetened beverages on appetite and energy intake in healthy adults following acute consumption but also after 3-month stevia consumption [3,4]. Understanding the mechanisms mediating the effects of NNS on appetite, food intake and health is of major nutritional and clinical importance, as their wider use could constitute a promising strategy for obesity management.

The gut microbiota is considered one of the key elements contributing to the regulation of host health [5]. Therefore, any changes in the composition or quality of the gut microbiome may have physiological consequences for the host. Host genetics, environmental factors and diet are considered the most important factors affecting microbiome composition [6,7]. There is a well-established relationship between certain microbial populations, insulin resistance and Type II Diabetes Mellitus, implying that the gut microbiome affects glucose regulation [8,9]. A potential link has been proposed to exist between sweetener consumption, glucose metabolism and gut microbiome composition. Suez *et al*. showed that the ingestion of saccharin by animals and humans induced alterations in metabolic pathways linked to glucose intolerance and dysbiosis in humans [10]. Despite this initial demonstration, more recent studies examining the effects of aspartame, sucralose and saccharin consumption on gut microbiota composition and included larger sample sizes and longer exposure duration, did not show any significant changes in the gut microbiome [11–13]. However, a further study by Suez et al. showed personalised effects of NNS on the microbiome [14]. To our knowledge, there are few other human trials that have assessed gut microbiome changes following repeated exposure to stevia, with a dose comparable to general consumption by the public.

Steviol glycosides, the sweet compounds of stevia, do not undergo degradation in the upper gastrointestinal tract, but they enter the colon intact and are degraded into glucose and steviol by the gut bacteria [15]. Steviol is absorbed and reaches the liver where it conjugates with glucuronic acid to facilitate secretion [16,17]. Therefore, stevia may have a brief point of contact with the gut microbiome. Whether this contact is enough to induce changes in the gut microflora composition remains to be investigated.

Using stored faecal samples from our previously published, randomized, controlled and open-label trial that showed an effect on body weight [3], the present study aimed to examine the effects of daily stevia consumption, in real-world quantities, on gut microbiota composition, diversity and community structure in healthy volunteers.

## Materials and Methods

### Study Design

The study was conducted as per detailed in Stamataki et al [3]. This study is a randomised controlled open-label 2-parallel-arm trial conducted in healthy adults with normal body mass index (BMI). Participant assignment was random; using an online tool (www.random.com) an independent person created a random sequence of zeros and ones (zero meant that the participant receives no treatment, one meant that the participant receives stevia) that was pre-stratified by gender. Ethical approval was granted by the University of Manchester Research Ethics Committee (2018-4812-7661); all subjects signed informed consent before participation in the study and received compensation for their time. The trial was registered at clinicaltrials.gov under the registration NCT03993418.

In this study participants were randomly allocated to either daily consumption of stevia (in liquid form), that was administered as 5 drops of a commercially available product (SweetLeaf Stevia Sweet Drops Clear, Sweetleaf®, Arizona, USA) in habitual beverages twice a day ideally before lunch and before dinner (total 10 drops, 5 drops of stevia corresponds to the sweetness of one teaspoon of table sugar), or in the control group where no changes in usual diet were required.

### Participants

The full participant criteria is detailed in Stamataki et al [3]. In brief, healthy adults with a normal BMI (18.5–24.9 kg/m^2^), aged 18–40 years old, who were non-habitual consumers of NNS (≤1 can of diet beverages per week or ≤1 sachet of NNS per week) and non-restrained eaters (restraint eating score in the Dutch Eating Behaviour Questionnaire (DEBQ) ≤ 3) were recruited. Other inclusion criteria were fasting blood glucose ≤ 6 mmol/L, stable weight for the last 12 months (±5 kg), willingness to comply with the study protocol, and no self-reported food allergy or intolerance to foods supplied during the study. Exclusion criteria were being on a diet or having ceased a diet in <4 weeks, following any special diets for weight maintenance, being vegetarian or vegan, having alcohol consumption of more than 14 units a week, more than 10 h of vigorous physical activities per week and/or planning to increase or decrease physical activity levels in the future, having ceased smoking in the last 6 months, and female participants who are or may be pregnant or currently lactating.

Participants who were eligible to participate completed an online screening questionnaire, and if inclusion criteria were satisfied, they were invited to a screening session on site following an overnight fast. Fasting blood glucose, weight and height were measured, and all details of the study were explained to participants. Eligible participants were invited to the full study and consented prior to participation.

A sample size calculation was conducted for the primary outcome, which was glucose response to an oral glucose tolerance test (OGTT), as described in our previous publication [3]. A total of 28 participants completed the study and a detailed flowchart is provided (Figure S1). Faecal samples were collected from 14 participants in the stevia group and 13 participants in the control group.

### Study protocol

Participants randomised to the stevia group were required to start consuming stevia (in liquid form) with their habitual drinks twice a day, while the control group received no intervention. During the intervention period participants were required to avoid all other NNS in beverages or foods, and they received training regarding the hidden sources of any NNS by a dietician. In addition, participants were asked to refrain from consuming any probiotic supplements. The stevia product administered in this study was selected based on its purity, as it contained only stevia leaf extract and water. Due to the nature of this study, blinding cannot be performed. Participants were aware of the trial arm to which they are randomized, the stevia arm or the control arm. Therefore, we are not eliminating cognitive factors which may occur when subjects are consuming stevia.

Participants attended 3 study visits, one at baseline, one at 6 weeks and the last one at 12 weeks. For visit week 0 and visit week 12 participants fasted overnight, for visit week 6 they had to abstain from foods and drinks for 2 hours prior to the scheduled time visit. All study procedures were conducted at the Neuroscience and Psychiatry Unit at the University of Manchester. A summary of the study schedule for assessment is presented (Table S1).

A detailed description of the outcomes presented in the original study such as the glucose and insulin response to the oral glucose tolerance test, weight, energy intake, blood pressure, and appetite questionnaires can be found in Stamataki *et al*. [3].

### Faecal sample collection

Participants were required to collect a faecal sample at home one day prior to the study visit. Faecal sample collection kits were provided to the participants. They were advised to collect samples from 3 random sites of the stool. After collection, participants were asked to keep the sample immediately in cold storage (−4°C) for up to 24 hours and bring it with them in the morning. Once samples were collected at our lab, they were stored at −80°C until further analysis. Although the study was open-label, the processing and analysis of faecal samples were conducted by a blinded-researcher.

### Isolation of genomic DNA and PCR

200 mg of stool was collected from each participant and microbial DNA extracted using a Qiagen DNeasy PowerSoil Pro Kit (Qiagen, Manchester, UK), in line with manufacturer’s instructions. First-round PCR was performed to amplify the V4 hypervariable region (V4_515fb.F: 5’-TCGTCGGCAGCGTCAGATGTGTATAAGAGACAGGTGYCAGCMGCCGCGGTAA-3’ and 4_806rB.R: 5-GTCTCGTGGGCTCGGAGATGTGTATAAGAGACAGGGACTACNVGGGTWTCTAAT-3’), using KAPA HiFi Hotstart ReadyMix (Roche, Burgess Hill, UK).

### 16S rRNA sequencing

54 samples, including additional experimental controls, were submitted for 16S rRNA sequencing. The Illumina 16S metagenomic Sequencing Library Preparation protocol was followed. All samples underwent amplicon PCR clean-up using AMPure beads (Beckman Coulter, Indianapolis, USA). Illumina sequencing adapters and dual index barcodes were added to each library using an Index PCR (Illumina XT Index Kit v2) followed by PCR clean-up with AMPure beads (Beckman Coulter). Libraries were quantified using the Qubit Fluorometer and the Qubit dsDNA HS Kit (ThermoFisher Scientific). Library fragment-length distributions were analyzed using the Agilent TapeStation 4200 and the Agilent D1000 ScreenTape Assay (Agilent Technologies, California, USA). Libraries were then pooled in equimolar amounts. Library pool quantification was performed using the KAPA Library Quantification Kit for Illumina (Roche). The library pool was sequenced on an Illumina MiSeq using a MiSeq v2 500 cycle Kit (Illumina; MS-102-2003) to generate 250-bp paired-end reads.

### Microbiome data analysis

Data was analysed using QIIME 2.0 [18], with DADA2 [19] as the de-noising step. The mean number of reads in experimental samples after quality filtering was 179,850 (range 93,060-426,047). There were 3 reads in a negative control of purified water after de-noising, which was deemed comparatively negligible. Microbiome data was processed using the Phyloseq R package [20], version 1.3.6. To avoid bias caused by sequencing depth during diversity analysis, data were rarefied to 61,353 reads (the lowest read depth). Beta diversity was calculated using weighted UniFrac [21] and plotted using principle coordinates analysis (PCoA) with the ‘plot_ordination’ function in the Phyloseq package. Jaccard Index values were also calculated amongst all samples and plotted as non-metric multidimensional scaling (NMDS) graphs using the ‘plot_ordination’ function, checking to ensure convergence in all cases. A scree plot was generated using the dimcheckMDS function in the ‘goeveg’ R package [22], version 0.5.1, to ensure that stress values were below the 0.2 acceptability threshold [23].

The relative abundance of each taxon was calculated at each taxonomic level. Taxa that made up less than 0.01% of the total abundance for phyla, class and order, less than 0.02% for family and 0.011% for genera, were merged into one group. Differential abundance between taxa was calculated using the DESEQ2 R package [24], version 1.32.

Random forest analysis was performed with the ‘randomForest’ R package, version 4.6, using relative abundances based on identified genera. ‘Out of Bag Error’ (OOB) values were obtained and used to estimate the predictive accuracy of the model in finding associations between the microbiome and the different treatment groups. To validate these associations, taxon abundance and samples were randomised to create a ‘negative control’ model, in order to see whether these associations were simply due to chance. Each forest was controlled for all other treatments (i.e. a random forest predicting treatment also included time as an explanatory variable). Taxa most important for making distinctions were identified based on their ‘MeanDecreaseAccuracy’ values.

### Statistical Analysis

Statistical differences between groups in beta diversity were calculated with PERMANOVA, using the adonis function the ‘vegan’ [25] R package, version 2.5.7. Differences in alpha diversity between groups was calculated using pairwise Wilcoxon Rank Sum tests using the ‘pairwise.wilcox.test’ function in R. Statistical analysis on differential abundance was performed with the calculation of geometric means before estimation of size factors, in the same software, using the DESeq2 package and the DESeq function. Significantly different taxa were defined as those with p < 0.01.

## Results

### No broad changes in the gut microbiota between treatments

To investigate large-scale patterns in the data, beta diversity was calculated using weighted UniFrac and plotted as a PCoA (Figure 1). It can be seen that separation along the x-axis explained 90.9% of the variation within the data, which visually appears to separate by stevia treatment. However, PERMANOVA suggested that there were no significant differences between any of the comparisons, combining all treatments (P = 0.947), comparing time i.e. 0 weeks vs 12 weeks (P = 0.121) and comparing treatment i.e. control vs stevia (P = 0.978). A similar analysis using the Jaccard Index also showed no differences between groups, comparing all treatments combined (P = 0.981), time (P = 0.9964) and treatment (P = 0.981) (Figure S2). The stress value was 0.17 (Figure S2), within the 0.20 acceptability threshold.

**Figure 1:**
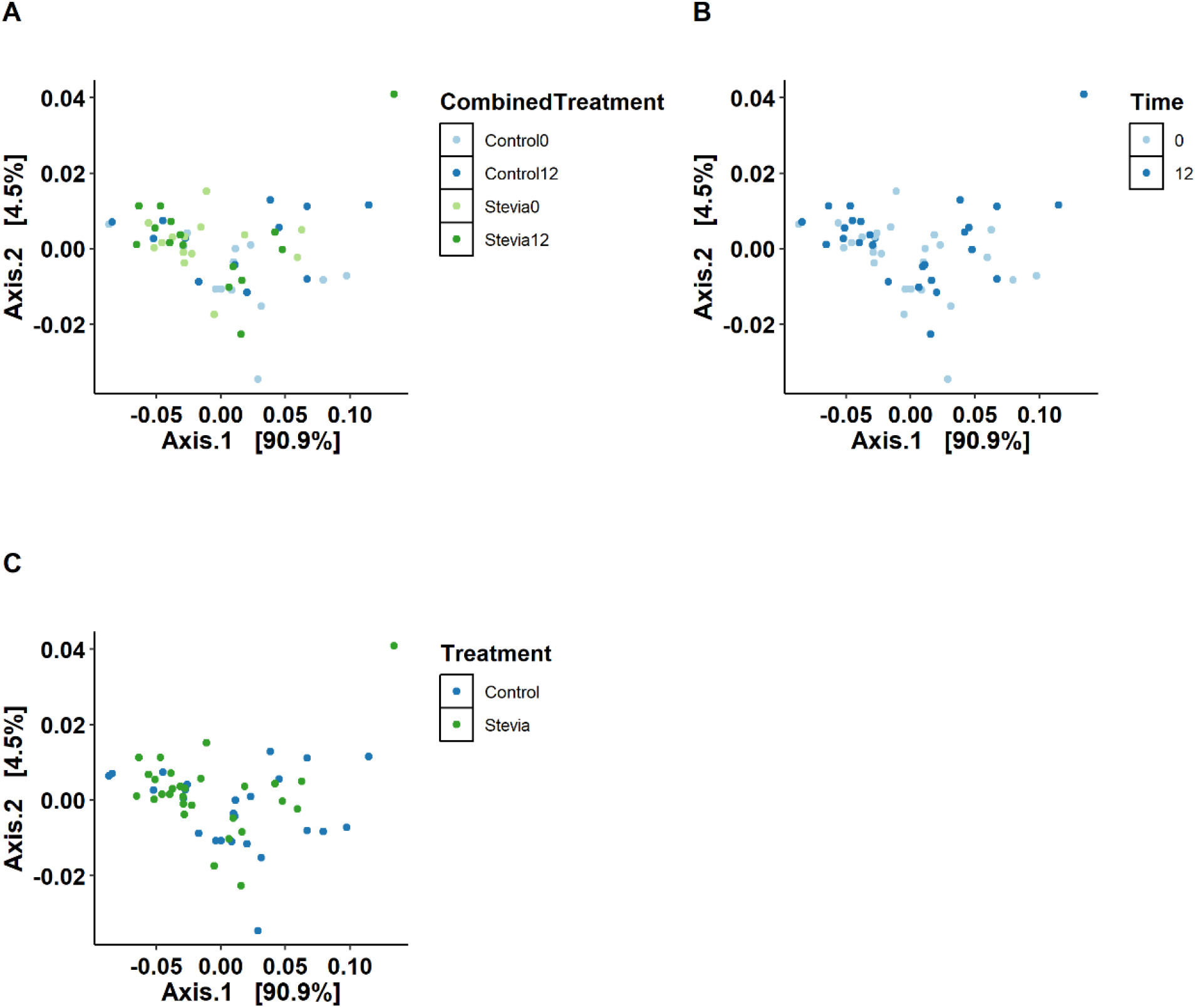
Principle Coordinates Analysis (PCoA) of gut microbiome data from control vs stevia groups. Healthy participants were asked to consume five drops of the sweetener stevia, twice daily, compared to control participants. Stool samples were collected from these participants at baseline (0 weeks) and 12 weeks after the intervention. Bacterial DNA was extracted from 200mg of stool and 16S rRNA sequencing was performed to analyse the gut microbiome. PCoA was plotted using the weighted UniFrac method, comparing all experimental groups (A), time only (B) and control vs stevia irrespective of time (C).

Alpha diversity was calculated to give a measure of sample richness (number of different taxa) and evenness (spread of taxa abundance) between groups (Figure 2). All groups had a median of around 300 observed OTUs, with no significant difference between groups (p = 0.966, when comparing all treatments). Analysis of evenness showed that all groups were comparable with relatively even community composition (median values > 0.75). There were no significant differences when comparing all treatments (p = 0.346) and there were similar levels of intra-group variability (Figure 2B). Analysis of the Shannon Index, which takes into account both evenness and richness, showed no significant difference between groups when comparing all treatments (p = 0.652) (Figure 2C). All groups had a large amount of variability.

**Figure 2:**
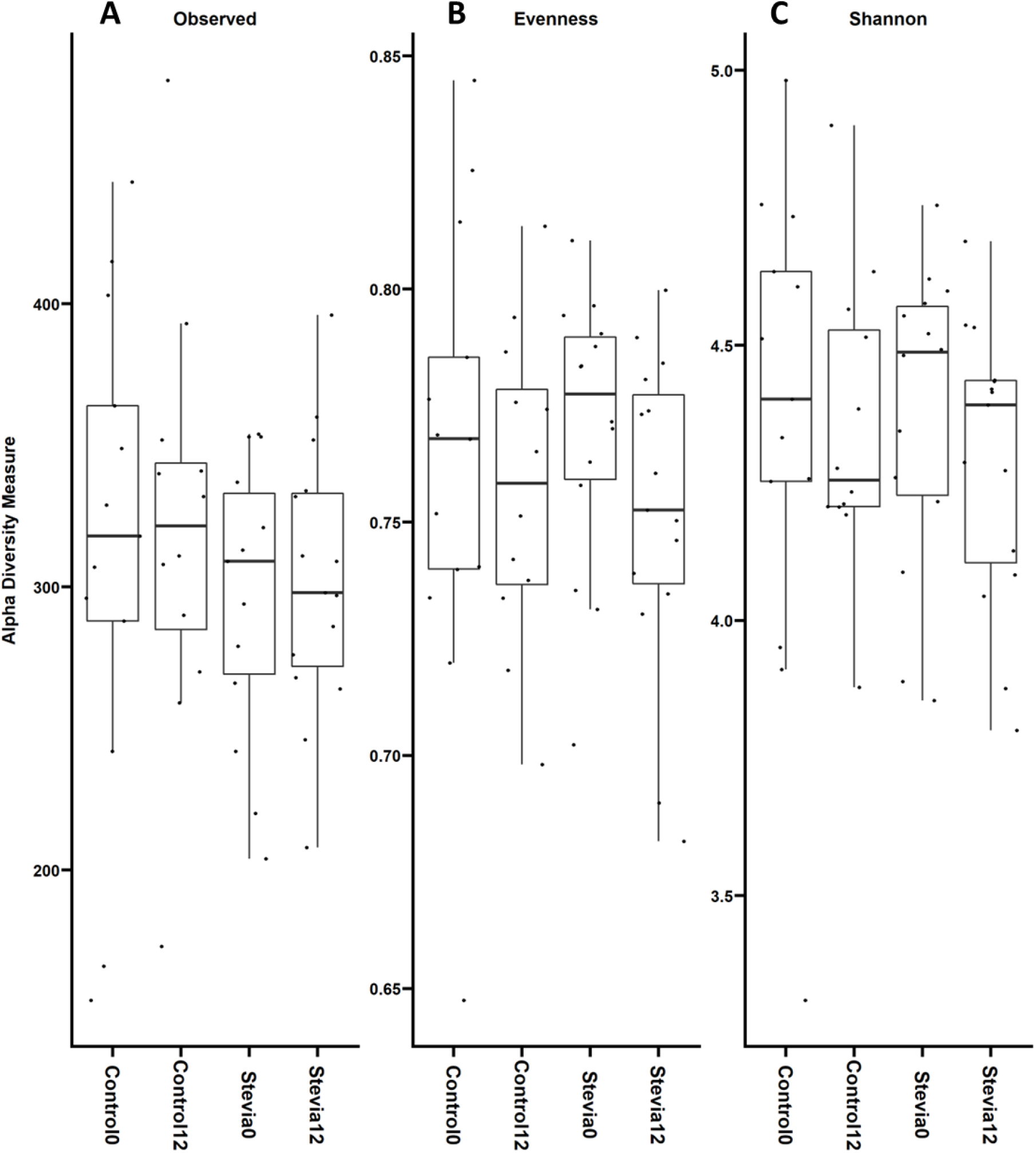
Alpha diversity analysis of gut microbiome data from control vs stevia groups. Healthy participants were asked to consume five drops of the sweetener stevia, twice daily, compared to control participants. Stool samples were collected from these participants at baseline (0 weeks) and 12 weeks after the intervention. Bacterial DNA was extracted from 200mg of stool and 16S rRNA sequencing was performed to analyse the gut microbiome. Alpha diversity, in terms of observed taxa, evenness and Shannon Index was plotted comparing all groups. Individual data points are also shown on each graph.

### Taxa abundance is similar between control and stevia groups

The taxonomic composition of the data comprised 15 phyla, 27 classes, 42 orders, 68 families and 130 genera (the numbers refer to unique taxa only). When taxa abundance was investigated at the phylum level (Figure 3A), the dominant phyla were Firmicutes, Bacteroidetes and Actinobacteria. At baseline (0 weeks) the control and stevia group had similar proportions of phyla, although the proportion of Actinobacteria appeared to be lower in the stevia group at both timepoints. Additionally, virtually no Proteobacteria in the stevia group could be visualised. At the genus level (Figure 3B), relative abundances were similar between the control and stevia groups. However, at 12 weeks, *Clostridium* and *Dorea* could be identified in the stevia group, which were not present at baseline. *Clostridium* and *Megamonas* were no longer present in the control group at 12 weeks, whereas *Dorea* could also be identified at 12 weeks in this group. Relative abundances at the class, order and family level were also plotted (Figure S3), although no major differences were visualised.

**Figure 3:**
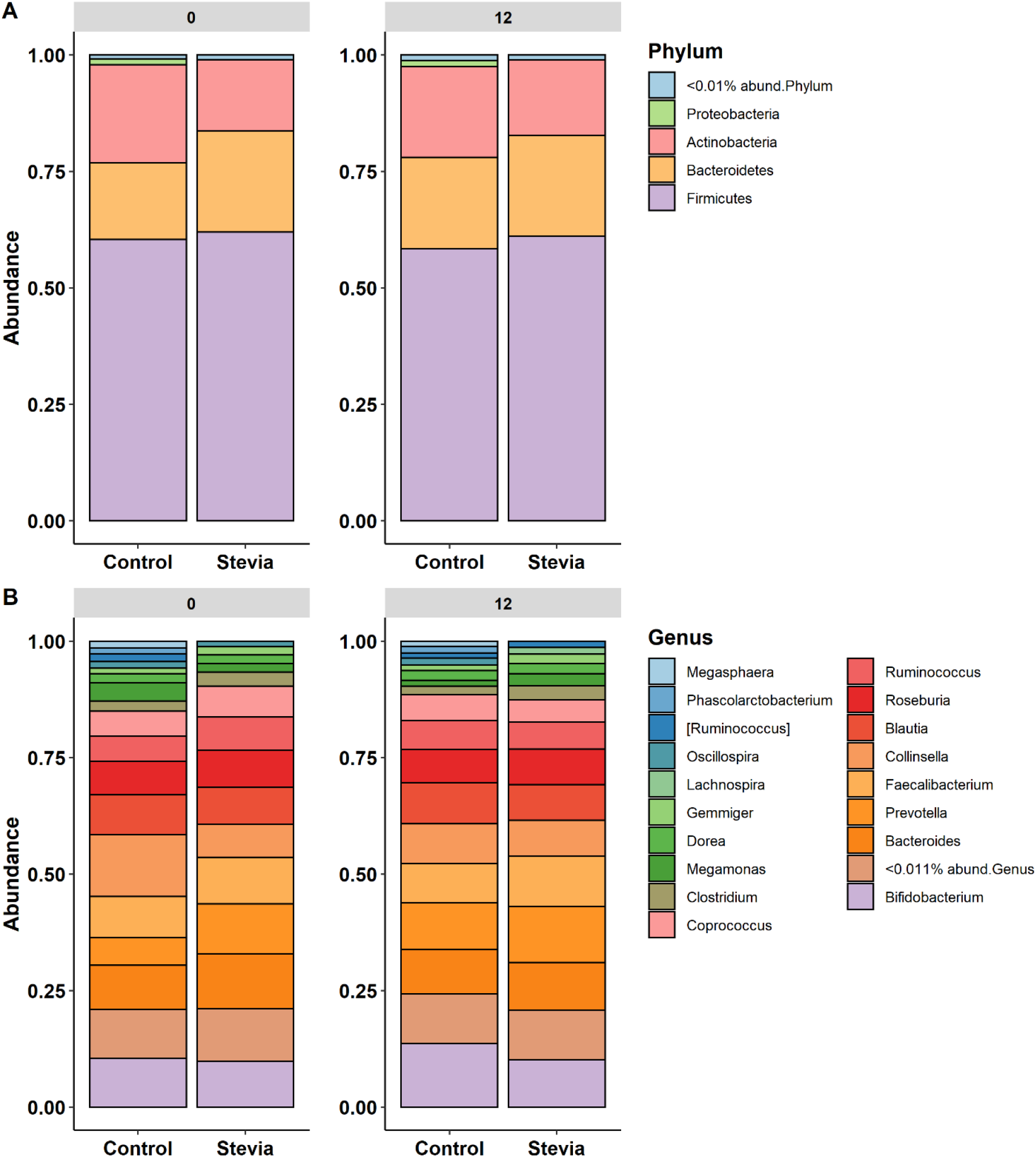
Taxa abundance compared between control vs stevia groups. Healthy participants were asked to consume five drops of the sweetener stevia, twice daily, compared to control participants. Stool samples were collected from these participants at baseline (0 weeks) and 12 weeks after the intervention. Bacterial DNA was extracted from 200mg of stool and 16S rRNA sequencing was performed to analyse the gut microbiome. Taxa abundance was plotted for phyla (A) and genera (B), with taxa less than 0.01% or 0.011% abundance respectively, combined into a single fraction.

Differential abundance was calculated to identify which taxa, if any, were statistically significantly different between groups. When the control groups were compared at baseline and 12 weeks, 15 genera were significantly different between the groups (Figure 4A). Variation also existed at baseline between the control and stevia groups (Figure 4B), with 19 significantly different genera. When stevia at baseline and at 12 weeks was compared (Figure 4C), there were only 9 significantly different genera between the two timepoints. Notably, stevia treatment led to a significant decrease in *Akkermansia* and an increase in *Faecalibacterium*. When control and stevia groups were compared at 12 weeks (Figure 4D), many of the significantly different taxa were already significantly different at baseline. *Butyricoccus* was the only genus identified as significantly different at 12 weeks that was not already different at baseline. However, some species were significantly different at baseline, but not after 12 weeks of stevia consumption. These bacteria included *Collinsella* and *Aldercreutzia*. Additionally, there were two *Coprococcus* species identified as significantly different at baseline (one higher and one lower when comparing stevia vs control). However, after 12 weeks, both species were significantly elevated in the stevia group.

**Figure 4:**
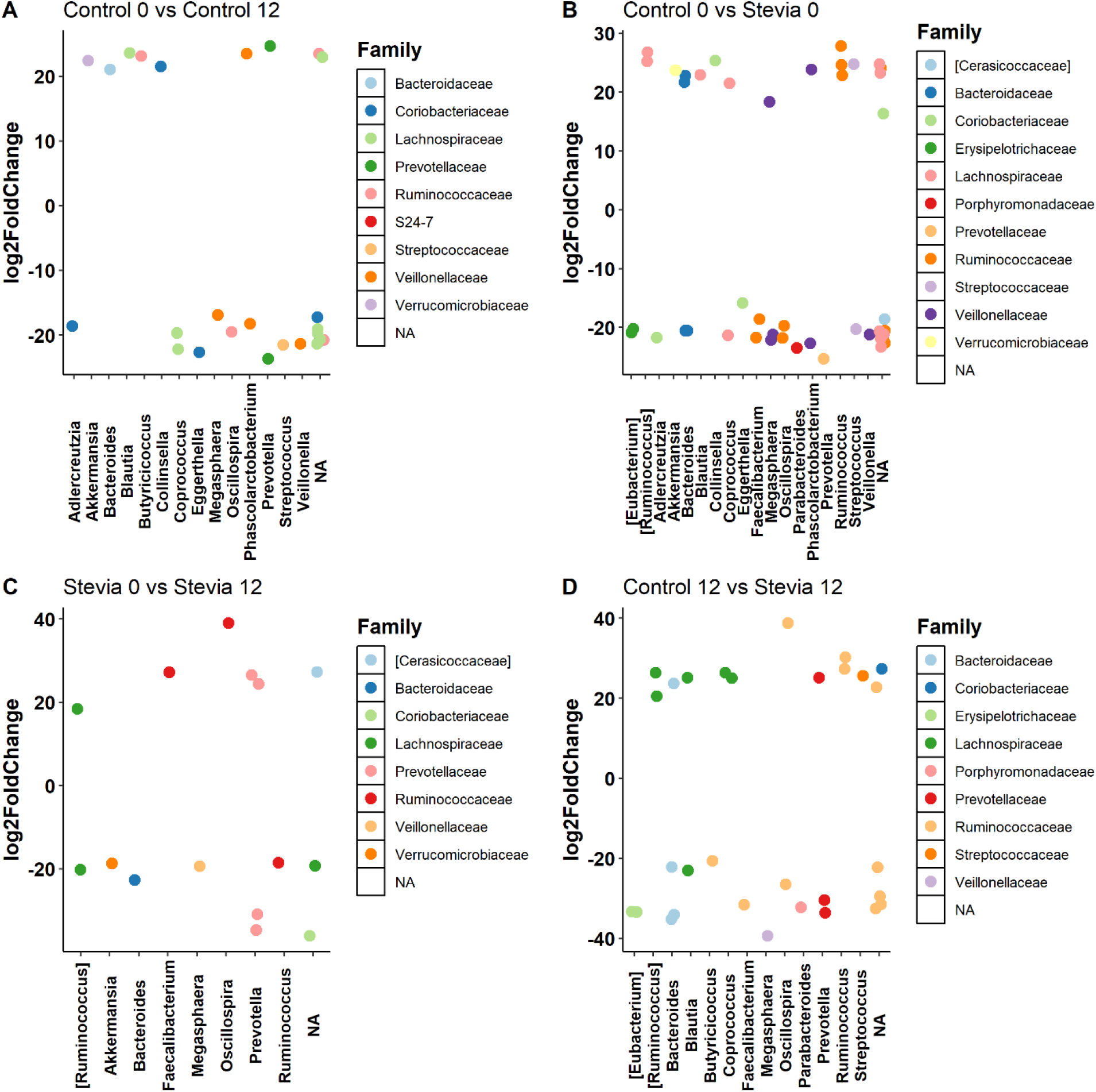
Differentially expressed bacteria between groups. Healthy participants were asked to consume five drops of the sweetener stevia, twice daily, compared to control participants. Stool samples were collected from these participants at baseline (0 weeks) and 12 weeks after the intervention. Bacterial DNA was extracted from 200mg of stool and 16S rRNA sequencing was performed to analyse the gut microbiome. The differential abundance of taxa between groups was calculated using the DESeq2 R package.

### Random forest finds associations between microbiome and stevia group

Random forest was employed to investigate any associations between the gut microbiome and the different groups in this study (Figure 5). Significant associations could be found between the gut microbiome when comparing control vs stevia (Figure 5A) (p < 0.05). Indeed, the model was able to associate the microbiome with each treatment correctly with approximately 75% accuracy. The negative control model had only ~50% accuracy. The taxa most strongly associated with this comparison were *Dehalobacterium, Methanobrevibacter, Oscillospira and Oxalobacter*. No significant associations could be found when comparing time (Supplementary Figure 4A) or when the treatment groups were combined (Supplementary Figure 4B). Taxa that were identified as most important for time are shown (Figure S5A). Notably, two genera were shared between the treatment only comparison and when treatment groups were combined, *Dehalobacterium* and *Oxalobacter* (Figure S5B).

**Figure 5:**
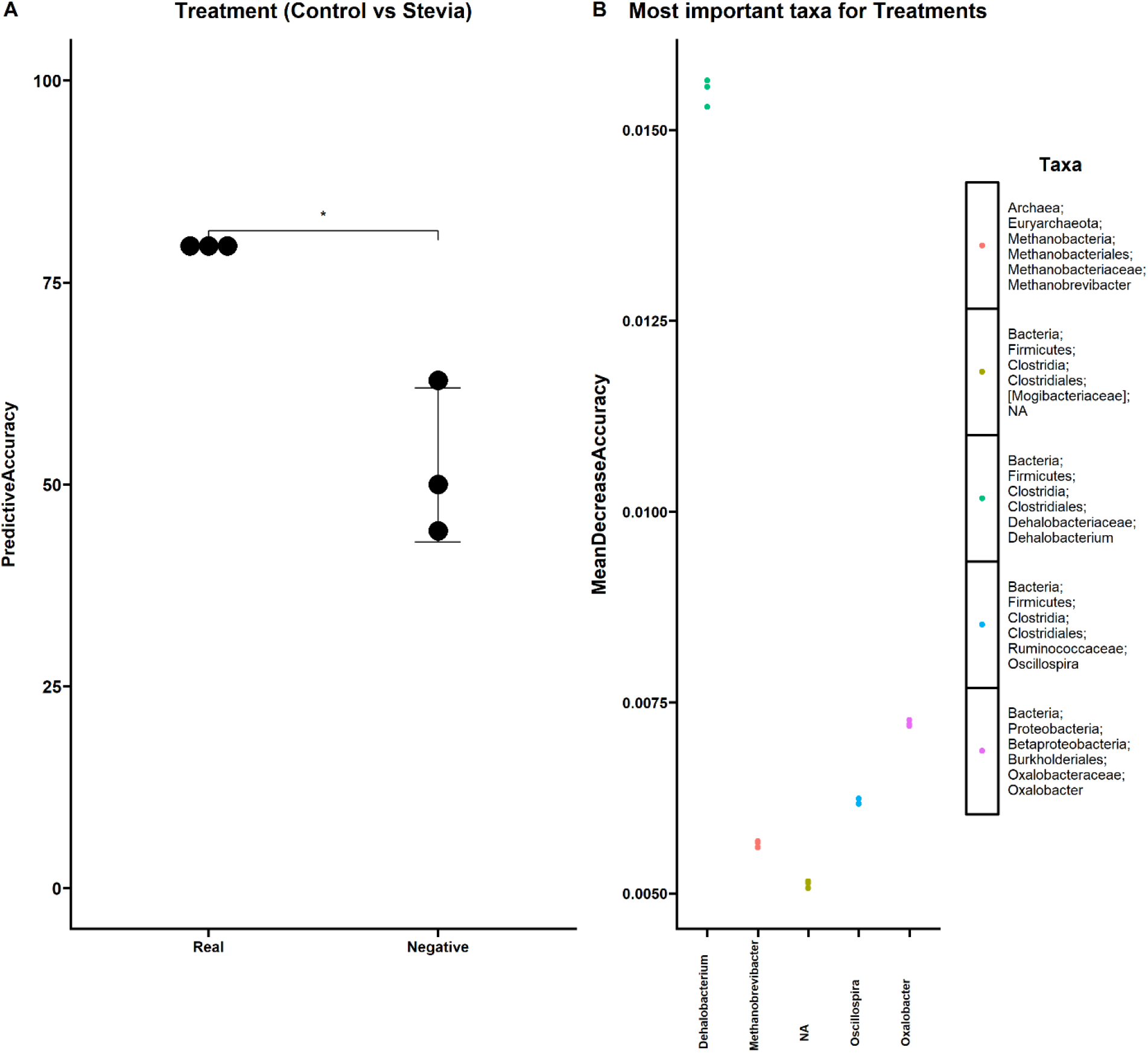
Associations between the gut microbiome and stevia group. Healthy participants were asked to consume five drops of the sweetener stevia, twice daily, compared to control participants. Stool samples were collected from these participants at baseline (0 weeks) and 12 weeks after the intervention. Bacterial DNA was extracted from 200mg of stool and 16S rRNA sequencing was performed to analyse the gut microbiome. Random forest was used to find associations between the relative abundance of identified genera and the accuracy of the model (A) and the genera most strongly associated with these groups (B). Statistical significance was determined via Student’s t-test (* p < 0.05). Individual data points are shown (+/- SEM).

## Discussion

We investigated how the use of stevia could impact the human gut microbiome in a randomised controlled open-label trial. Various studies have investigated the impact of stevia in different contexts such as mice [26] or using *in vitro* models [27], but to our knowledge, few clinical studies have specifically investigated the impact of stevia on the gut microbiome. Some clinical studies investigated the use of alternative NNS, such as saccharin [10,13], sucralose [11] and aspartame [11,28]. A key initial study that investigated the impact of NNS on the microbiome was the study by Suez et al. [10], who found that saccharin consumption led to glucose intolerance induced by microbial gut dysbiosis. This study was followed by a comprehensive assessment of various other NSS, including stevia, where they found that sucralose and saccharin impaired the glycaemic response [14]. However, all NSS tested had a significant impact on microbiome function [14]. In comparison, their study was conducted over a shorter time (2 weeks), compared to 12 weeks in our present study. We previously found no difference in glucose tolerance following long term stevia consumption [3] and in our present study, we found no significant differences between the treatment groups when comparing beta and alpha diversity (Figures 1–2). Different sweeteners have different chemical compositions (summarised by Turner et al. [29]) and therefore, their functional impacts on the host and their metabolism by the microbiota are likely to be different.

However, our findings are similar to the study by Ahmad et al. [11], who found that treatment with aspartame or sucralose in healthy adults had no discernible impact on these metrics. These findings are in contrast to another study [28], which showed significant differences in bacterial diversity after only 4 days of aspartame or acesulfame consumption. Although we do not see significant differences in diversity, our PCoA plots showed strong separation along the x axis (Figure 1A-C), which can also be seen visually in how stevia and control participants separate on the graph. While these data may imply baseline differences in the community composition between control and stevia participants, it was not a significant difference. Notably, all groups had comparable alpha diversity, suggesting that community composition in each group was relatively even over time and equally diverse.

Changes in the gut microbiome, at any taxonomic level, are often associated with functional impacts on the host [30,31]. In our study, at the phylum level, the taxonomic profile closely resembled a typical gut profile (Figure 3A), dominated by Firmicutes and Bacteroidetes, but Actinobacteria were also prevalent, with a smaller fraction of Proteobacteria and other phyla. The ratio of Firmicutes to Bacteroidetes has often been explored in the context of obesity [32,33] and indeed, the use of NNS is frequently associated with energy intake and weight [1,34]. As discussed, our previous study investigated the impact of stevia on glucose tolerance, but body weight and energy intake were also explored [3]. We found that although there were no differences in glucose tolerance, there were significant differences in energy intake after 12 weeks of stevia consumption [3]. Based on our current observations, the impact on energy intake is unlikely to be mediated by the Firmicutes to Bacteroidetes ratio between control and stevia participants. The microbiome is strongly associated with the production of short-chained fatty acids (SCFAs) that may mediate relevant functional effects on the host [35,36]. One aspect that could be investigated in our future work is the production of SCFAs in response to stevia-treatment, alongside microbial profiling.

We show relative abundance at each taxonomic level for phyla and genera (Figure 3) and at the class, order and family level (Figure S3). Differential abundance revealed that only one taxon identified was significantly different between control and stevia groups after 12 weeks, *Butyricoccus*. This was downregulated after the consumption of stevia. One study investigated the impact of various sweeteners in rats and found that honey downregulated *Butyricoccus pullicaecorum* [37], a butyrate-producing species [38]. Blautia was significantly upregulated at baseline and significantly downregulated after 12 weeks of stevia consumption. In contrast, an *in vitro* study revealed that stevia treatment of human faecal samples led to no significant differences in the expression of *Blautia coccoides* [27]. Blautia is associated with the production of butyric and acetic acid [39]. Thus, it could be that stevia is leading to a reduction in butyrate-producing species. Indeed, after 12 weeks compared to baseline, the stevia group had a significant reduction in *Megasphaera* bacteria, with some species associated with butyrate production [40]. However, this was not significantly different when compared to the control group.

Many of the bacteria highlighted by the differential abundance analysis tend to be those identified in studies investigating weight and metabolism. For instance, one study found that *Blautia* was inversely correlated with the accumulation of visceral fat [41]. Additionally, we found that *Collinsella* was significantly higher at baseline, but no longer after 12 weeks of stevia treatment. Low dietary fibre intake has been shown to increase this genus in overweight pregnant women [42]. In context, we showed that participants consuming stevia significantly maintained their weight compared to control participants [3].

Our random forest identified taxa that were strongly associated with either control or stevia participants (Figure 5), that were not identified by differential abundance analysis. Notably, the most important microorganism identified was *Dehalobacterium*, a bacterium previously associated with a high BMI [43]. A study in mice revealed that sucralose treatment had a notable impact on the family *Dehalobacteriaceae* [44]. Curiously, these bacteria were also found in significant abundance in the intestinal mucus of a colitic-mouse model [45]. Previous studies have highlighted the difference between stool and mucus-resident bacteria [30,46] and therefore, there could be changes in the mucus bacteria that we do not see in this study.

Among the strengths of our study, is that presently, this is one of a limited number of studies to investigate the effects of stevia consumption on the human gut microbiome, where the dose of stevia reflected a normal dose to simulate regular consumption by the general population. The stevia used was also a pure stevia product (stevia in water), so any effects demonstrated can be linked to stevia. However, some limitations should be reported. The study was powered for the primary outcome which was postprandial glucose response to an oral glucose tolerance test [3], gut microbiome analysis was an exploratory secondary outcome as reported in the study registration. Additionally, as discussed, more functional analysis with respect to metabolite analysis could further strengthen this study.

Overall, although there may be several individual taxa that are associated with stevia use, we found no significant differences in overall community composition after 12 weeks of stevia consumption. Therefore, our data suggest that the regular, long-term consumption of stevia does not significantly impact upon the human gut microbiome.

## Supplementary Material

A flow chart summarising the clinical study is provided (Figure S1: Participant flow chat). We have also included supplementary data pertinent to ordination (Figure S2: NMDS of gut microbiome data plotted using Jaccard Index), relative abundance (Figure S3: Relative taxa abundance compared between control vs stevia groups) and our random forest models (Figure S4: Associations between gut microbiome and stevia group; Figure S5: Important taxa identified for random forest model). We also include a table summarising assessment criteria and procedures (Table S1: Summary of the study procedures and assessment).

## Acknowledgements

The authors would like to thank the contributions of the core facilities at the University of Nottingham, particularly Victoria Wright and Matt Carlile at the DeepSeq facility.

## Author Contributions

GS prepared samples for sequencing, performed the microbiome analysis, made figures and co-wrote the manuscript. AJM supervised the microbiome aspect of the project and critically reviewed and edited the manuscript. JTM designed and supervised the clinical aspect of the project and critically reviewed and edited the manuscript. NSS designed and facilitated the clinical study, made figures and tables and co-wrote the manuscript.

## Data Availability

Sequence data analysed during the current study will be made available. Code to reproduce the main text figures produced in R will be available on FigShare.

