## Supplementary Information for "Consumption of the non-nutritive sweetener stevia for 12 weeks does not alter the composition of the gut microbiota"

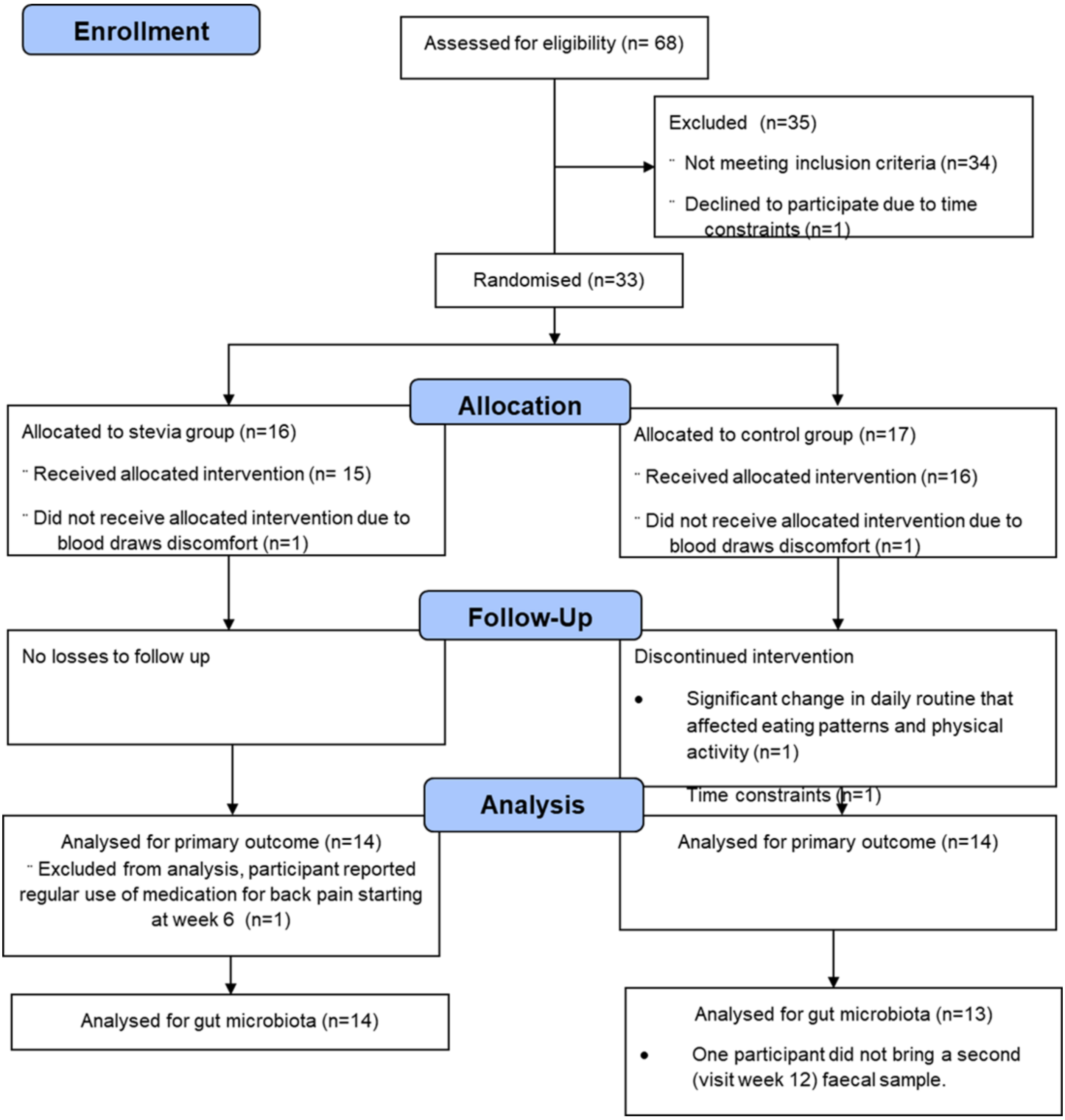


**Figure S1: Participant flow chat.**

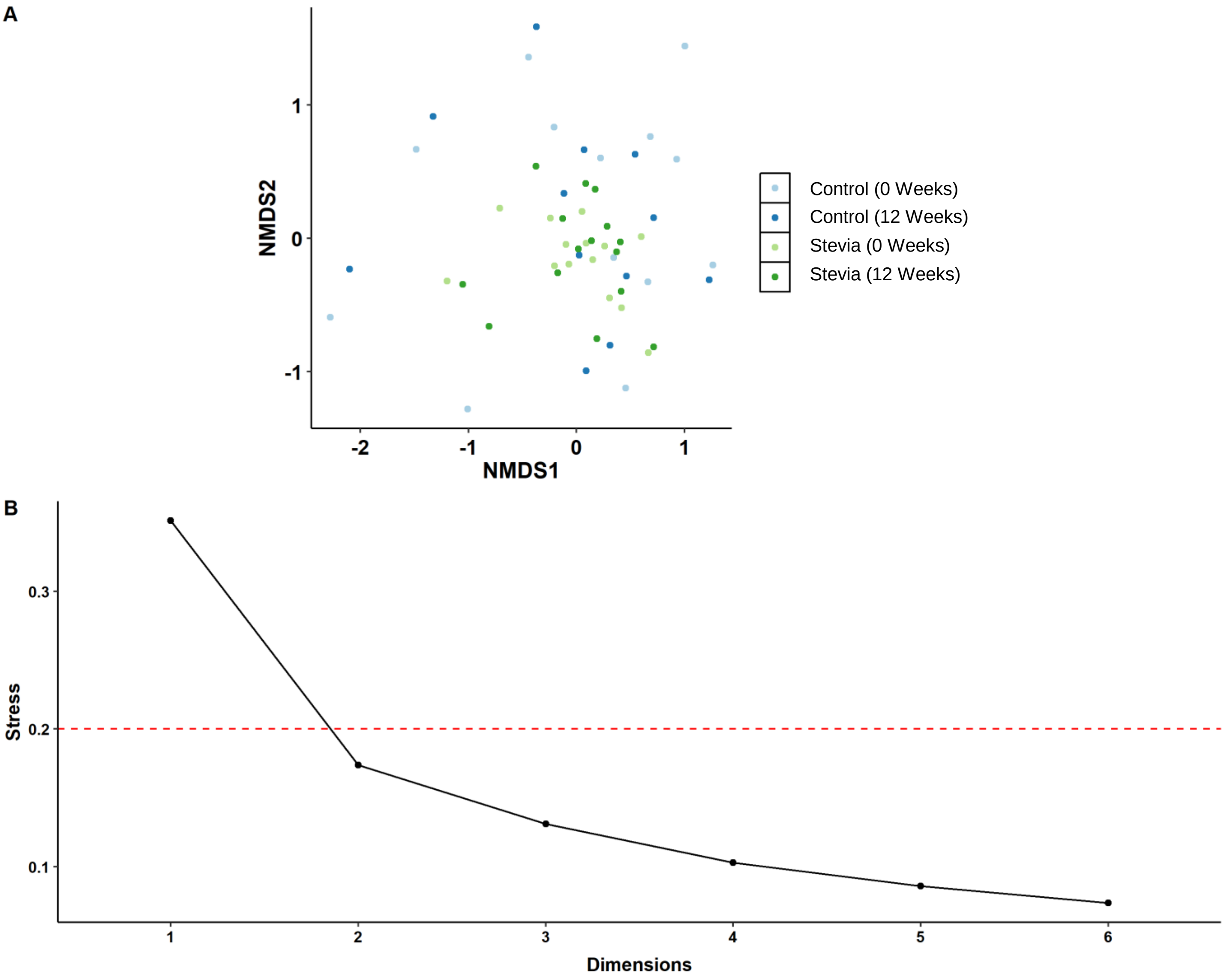


**Figure S2: NMDS of gut microbiome data plotted using Jaccard Index.** Healthy participants were asked to consume five drops of the sweetener stevia, twice daily, compared against control participants. Stool samples were collected from these participants at baseline (0 weeks) and 12 weeks after the intervention. Bacterial DNA was extracted from 200mg of stool and 16S rRNA sequencing was performed to analyse the gut microbiome. NMDS was plotted using Jaccard's Index (A). A scree plot was used to confirm the stress value was within the 0.2 acceptability threshold (B).

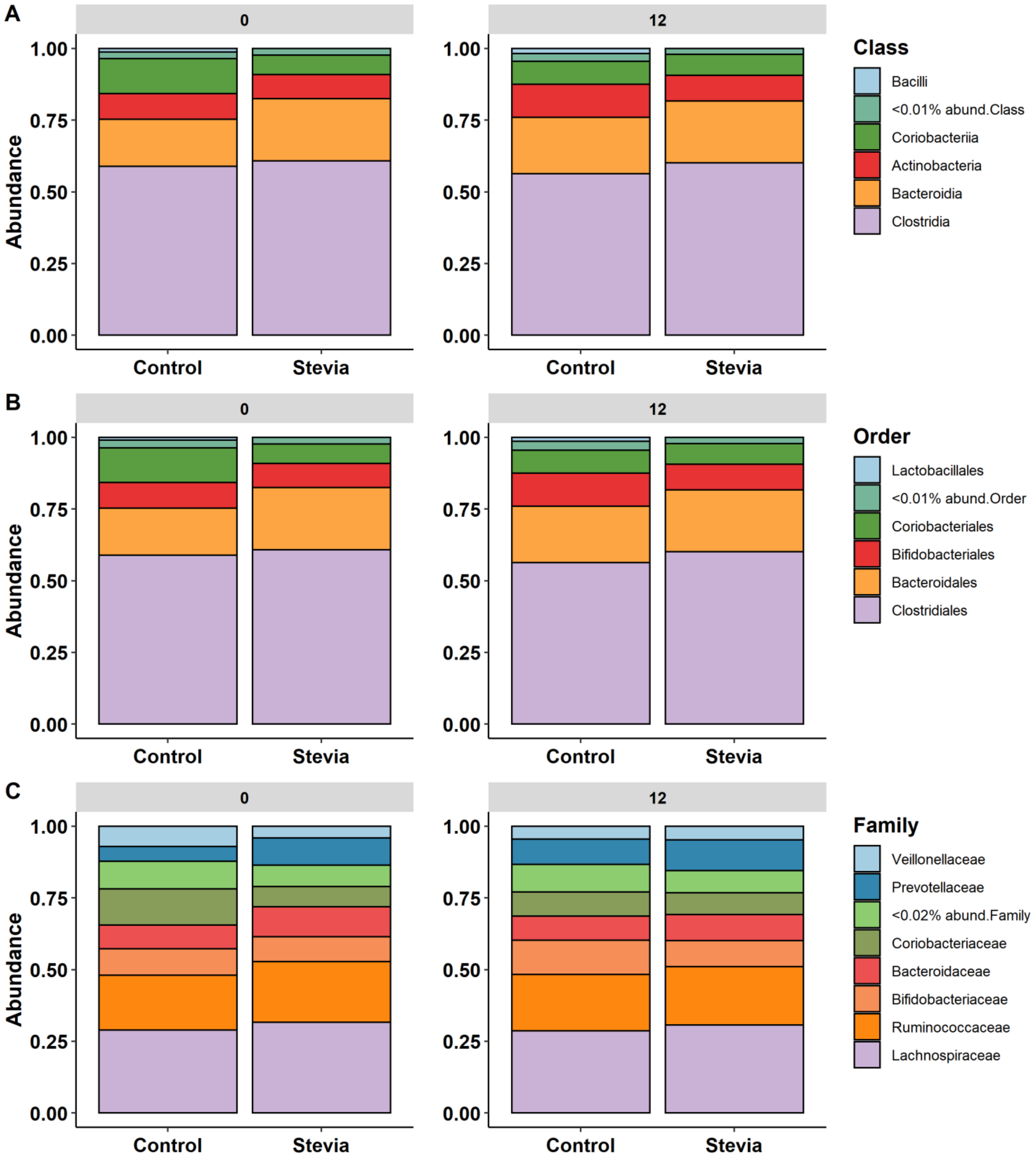


**Figure S3: Relative taxa abundance compared between control vs stevia groups.** Healthy participants were asked to consume five drops of the sweetener stevia, twice daily, compared against control participants. Stool samples were collected from these participants at baseline (0 weeks) and 12 weeks after the intervention. Bacterial DNA was extracted from 200mg of stool and 16S rRNA sequencing was performed to analyse the gut microbiome. Taxa abundance was plotted for class (A), order (B) and family (C), with taxa less than 0.01% abundance for class and order or 0.02% abundance for family, combined into a single fraction.

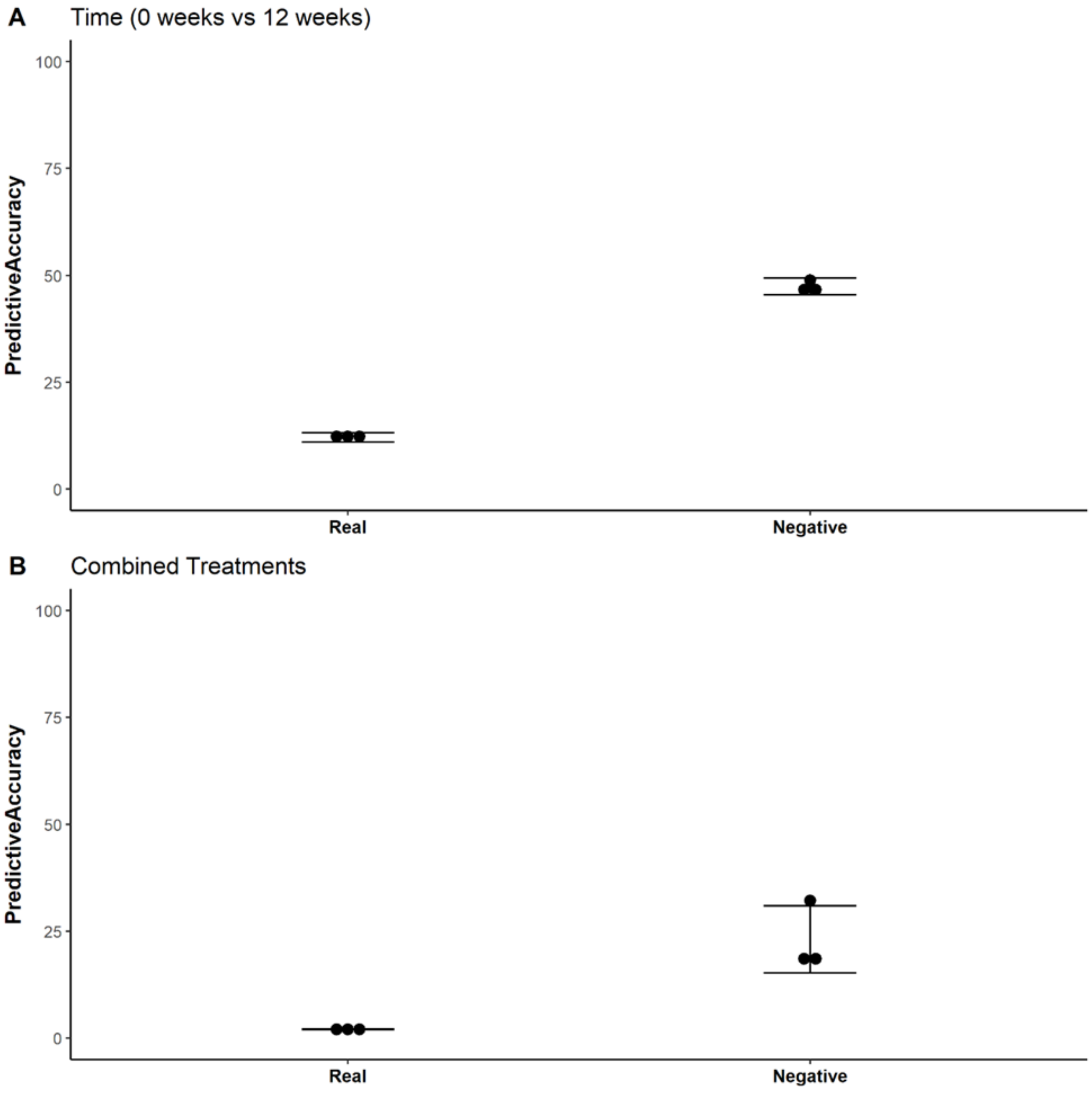


**Figure S4: Associations between gut microbiome and stevia group.** Healthy participants were asked to consume five drops of the sweetener stevia, twice daily, compared against control participants. Stool samples were collected from these participants at baseline (0 weeks) and 12 weeks after the intervention. Bacterial DNA was extracted from 200mg of stool and 16S rRNA sequencing was performed to analyse the gut microbiome. Random forest was used to find associations between the relative abundance of identified genera and the accuracy of the model for time (A) and combined treatments (B).

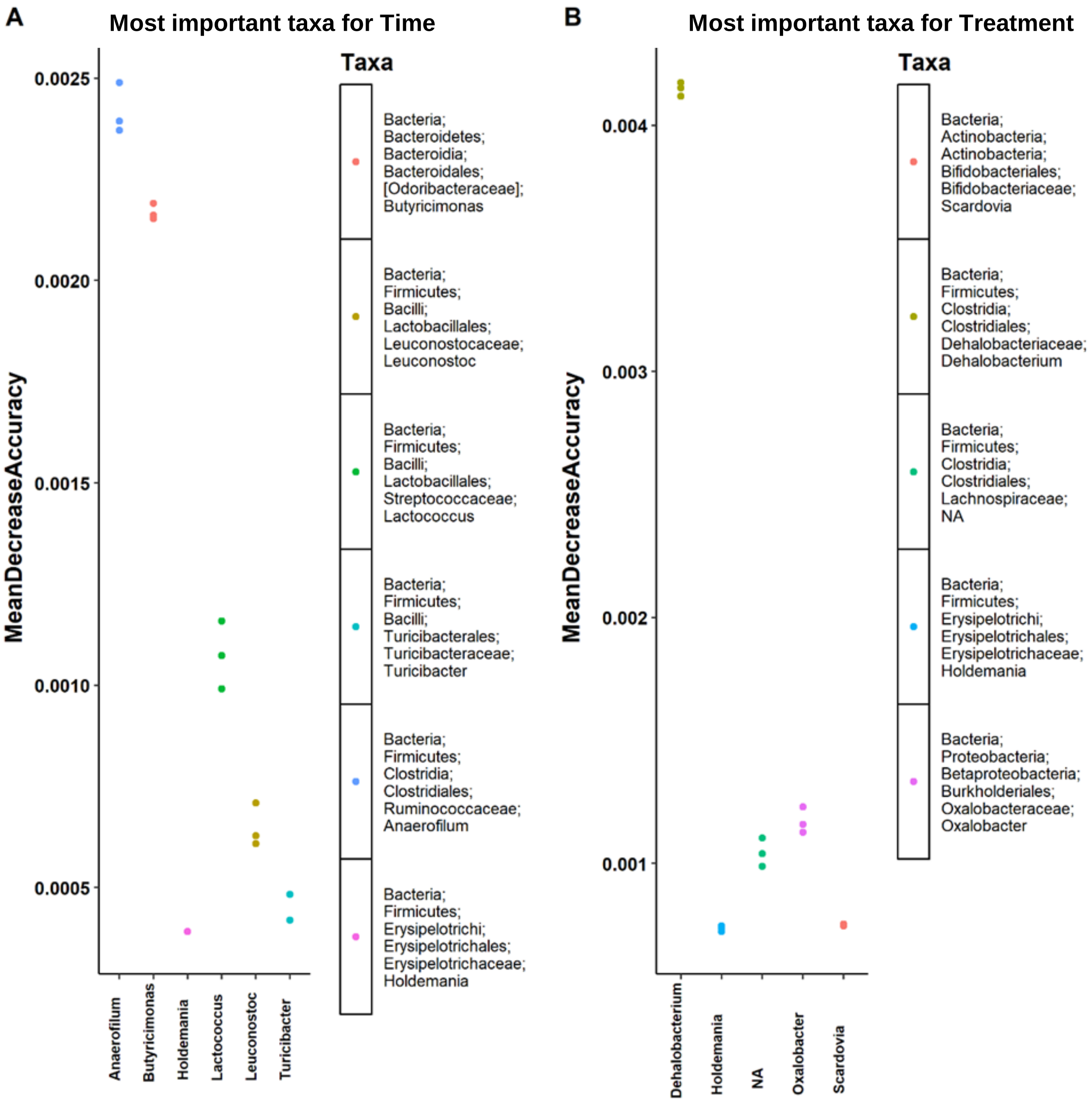


**Figure S5: Important taxa identified for random forest model.** Healthy participants were asked to consume five drops of the sweetener stevia, twice daily, compared against control participants. Stool samples were collected from these participants at baseline (0 weeks) and 12 weeks after the intervention. Bacterial DNA was extracted from 200mg of stool and 16S rRNA sequencing was performed to analyse the gut microbiome. Random forest was used to find associations between the relative abundance of identified genera and the accuracy of the model. The most important taxa associated with time (A) and combined treatments (B) are illustrated.

### Table S1: Summary of the study procedures and assessment.

| | Screening visit | Visit week 0 | Visit week 6 | Visit week 12 |
|---|---|---|---|---|
| General information, medical history | √ | | | |
| Informed consent | √ | √ | | |
| Weight | √ | √ | √ | √ |
| Height | √ | | | |
| Fasting blood glucose | √ | √ | √ | √ |
| Waist circumference | | √ | √ | √ |
| Oral glucose tolerance test | | √ | √ | √ |
| Blood draw session | | √ | | √ |
| 3-day dietary recall (Intake24) | | √ | √ | √ |
| International Physical Activity questionnaire | | √ | √ | √ |
| Blood pressure measurement | | √ | √ | √ |
| Appetite questionnaires | | √ | √ | √ |
| Faecal sample collection | | √ | | √ |
| Laboratory measurements in stool (faecal microbiome) | | √ | | √ |
